# *In vitro* comparison of SARS-CoV-2 variants

**DOI:** 10.1101/2023.03.11.532212

**Authors:** Kruttika S. Phadke, Nathaniel B. A. Higdon, Bryan H. Bellaire

**Affiliations:** Veterinary Microbiology and Preventive Medicine, Iowa State University, Ames, 50011; Immunobiology Interdepartmental Graduate Program, Iowa State University, Ames, 50011; Interdepartmental Microbiology Graduate Program, Iowa State University, Ames, 50011

**Keywords:** SARS-CoV-2, Vero E6, Calu3, A549, Cytopathic Effect (CPE), Viral replication

## Abstract

The Coronaviridae family hosts various coronaviruses responsible for many diseases, from the common cold, severe lung infections to pneumonia. SARS-CoV-2 was discovered to be the etiologic agent of the Coronavirus pandemic, and numerous basic and applied laboratory techniques were utilized in virus culture and examination of the disease. Understanding the replication kinetics and characterizing the virus’ effect on different cell lines is crucial for developing *in vitro* studies. With the emergence of multiple variants of SARS-CoV-2, a comparison between their infectivity and replication in common cell lines will give us a clear understanding of the characteristic differences in pathogenicity. In this study, we compared the cytopathic effect (CPE) and replication of Wild Type (WT), Omicron (B.1.1.529), and Delta (B.1.617.2) variants on 5 different cell lines; VeroE6, VeroE6 expressing high endogenous ACE2, VeroE6 highly expressing human ACE2 (VeroE6/ACE2) and TMPRSS2 (VeroE6/hACE2/ TMPRSS2), Calu3 cells highly expressing human ACE2 and A549 cells. All 3 VeroE6 cell lines were susceptible to WT strain, where CPE and replication were observed. Along with being susceptible to Wild type, VeroE6/hACE2/TMPRSS2 cells were susceptible to both omicron and delta strains, whereas VeroE6/ACE2 cells were only susceptible to omicron in a dose-dependent manner. No CPE was observed in both human lung cell lines, A549 and Calu3/hACE2, but Wild type and omicron replication was observed. As SAR-CoV-2 continues to evolve, this data will benefit researchers in experimental planning, viral pathogenicity analysis, and providing a baseline for testing future variants.

## 1. INTRODUCTION

Coronaviruses from the Coronaviridae family, are enveloped RNA viruses that can infect humans and animals (1). The most recently discovered coronavirus, SARS-CoV-2, was first identified in Wuhan, China, in late 2019 and was declared a pandemic in 2020 by the World Health Organization (WHO) (2,3). The disease COVID-19, caused by SARS-CoV-2, is estimated to be responsible for almost 700 million cases and 7 million deaths (4). Primarily a respiratory virus, it is transmitted mainly through aerosols causing symptoms like fever, cough, and fatigue, with more severe cases causing pneumonia and respiratory distress (5–7).

Throughout the pandemic, many variants of SARS-CoV-2 have emerged from different regions of the world that have quickly become a cause of concern. Two of these variants are delta (B.1.617.2) and omicron (B.1.1.529). Delta variant was first identified in late 2020 and is currently on the Variants being monitored (VBM) list by the Centers for Disease Control and Prevention (CDC)(8). It consists of almost 30 mutations which contribute towards increased infectivity and transmissibility, making it more dangerous than the Wild Type (WT) (9,10). Omicron variant was first identified in November 2021 and is currently the variant of concern (VOC) by the CDC (8). After sequencing, it was identified that this variant has almost 50 mutations, most of which are in the receptor binding domain of the spike protein. Due to its high transmissibility and ability to evade neutralizing antibodies, it rapidly made its way around the globe (11).

During investigations into the pathogenesis of coronavirus variants, it became increasingly difficult to compare results with SARS-CoV-2 variants, given their specific dependence on host cell types for replication. The lower cellular infectivity of recent variants confounds such a comparison since they require stable cell lines overexpressing cell receptors for routine culture. As a result, using different cell lines complicates the analysis of cellular assays, such as relative infectivity, virus replication rates, and antibody neutralization changes between WT and variant viruses. A relative comparison of variants across different cell lines assessing in vitro characteristics, such as cellular infectivity and sensitivity to neutralizing antibodies, would be monumentally beneficial to address such difficulties.

A benchmark permissive cell line for many viruses, including members of the coronaviridae, is the epithelial-like, African green monkey kidney cell line known as Vero E6, subcultured from original Vero cells in 1979. Vero E6 cells continue to be used for in vitro analysis of SARS-CoV-2, viral propagation, viral pathogenesis, pharmacology, diagnostics, and immunology research. The inability of Vero E6 cells to produce interferon following viral infection and the stable, higher expression of Angiotensin Converting Enzyme 2 (ACE2), the cell surface receptor for SARS-CoV-2 entry, compared to the parent Vero cell line make Vero E6 the preferred cell line for plaque-based assays (12–16). For *in vitro* studies such as antiviral testing, live virus neutralization assays and cytopathic effect (CPE) based assays, understanding the infection kinetics of the variants in the VeroE6 cell line is essential. Once the virus spike protein binds to ACE2 on the cell’s surface, the spike protein is cleaved by serine protease TMPRSS2 to initiate viral entry (15,17). Enhancing the expression of both ACE2 and TMPRSS2 can give us a better understanding of their importance in the viral infection cycle of the variants. We report our observations on the relative infectivity and replication of the Wild Type, Omicron, and Delta variants in multiple cell lines to identify variant specific characteristics. Here we have compared all three variants in VeroE6 cells, VeroE6 cells expressing high endogenous ACE2 and VeroE6 cells that are overexpressing human ACE2 and TMPRSS2. Lastly, the variants are also tested in the more physiologically relevant human lung cell lines, Calu3 overexpressing human ACE2 and A549 cells.

## 2. METHODS

### 2.1. Biohazard Statement

All infection experiments involving SARS-CoV-2 were performed in BSL-3 laboratory facilities at Iowa State University (ISU) with Institution Biosafety Committee approved protocols.

### 2.2. Cell culture, Virus strains, and virus amplification

African Green Monkey tissue culture cell line, Vero E6 (ATCC CRL-1586) cells and human lung epithelial, A549 cells (ATCC CCL185) were acquired from ATCC (Manassas, VA). VeroE6 high expressing ACE2 cells (NR53726) (VeroE6/ACE2), VeroE6 expressing TMPRSS2 and human ACE2 (NR54970) (VeroE6/hACE2/TMPRSS2) and Calu3 high expressing human ACE2 (NR55340) (Calu3/hACE2) were acquired from BEI (Manassas, VA). All cells were maintained in Dulbecco’s modified Eagle medium (DMEM) with 4.5g/L glucose, L-glutamine and sodium pyruvate (10-013-CV Corning) containing 10% fetal bovine serum (FBS) (Cytiva). Wild type (WT) SARS-CoV-2 (NR-52281) isolated from a COVID-19 patient in Washington, USA, Delta strain B.1.617.2 (NR-55671) and Omicron strain B.1.1.529 (NR-56481) were acquired from BEI. Viral infectious doses were prepared from supplied stocks by amplifying the virus through three passages, at which time aliquots were prepared and frozen. The stocks were quantified by Spearman and Karber algorithm (18). Wild-type virus was amplified with Vero E6 cells and Omicron and Delta variants were amplified using hACE2/TMPRSS2 overexpressing VeroE6 cell line.

### 2.3. Experimental infections

Cell cultures were seeded at a density of 2 x 10^4^ cells per well in 96-well plates 18 hours prior to infection. Viral stocks were thawed and diluted serially from 1000 Plaque forming units (PFU) to 0.01 PFU in DMEM with 5%FBS. Infection was initiated by replacing media on the cells with 100μL DMEM with 5%FBS per well and adding 100μL of each dilution to the cells. Each infection dilution was performed in triplicate. Infected cells were incubated for 5 days at 37°C under 5% CO2 atmosphere and observed under an inverted microscope every day. On day 5, the lactate dehydrogenase (LDH) enzyme levels were measured from 50μL of each cell culture supernatant using the CyQuant™ LDH Cytotoxicity Assay (Invitrogen C20301) according to the manufacturer’s protocol. Cell viability was calculated using the LDH released into the supernatant from infected cells at 490nm and 680nm. Cell viability was calculated using the equation,

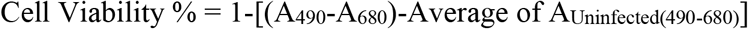

### 2.4. RNA isolation and RT-qPCR analysis

Viral genome equivalents were quantified from RNA harvested from cell culture supernatants. TRIzol reagent (Sigma) was used to extract RNA by mixing 400μL of TRIzol reagent with 100μL of cell culture supernatant. Nucleic acids were separated by adding 80μL of chloroform followed by centrifugation at 12000g at 4 °C for 15 minutes. RNA was precipitated by adding 200μL of isopropanol to the upper aqueous layer and centrifuging for 10 minutes at 12000g and 4°C. RNA pellet was washed with ethanol and resuspended in 50μL of nuclease-free water. RNA concentrations were normalized to 50ng/μL. RT-qPCR of the extracted RNA was carried out using Luna^®^ Universal Probe One-Step RT-qPCR Kit and IDT 2019-nCoV RUO Kit on the Bio-Rad CFX96 Real-Time system.

### 2.5. Statistical analysis

Data analysis, including calculations of average of means, standard deviation, or standard error and two-way ANOVA with a Bonferroni’s comparison of means was calculated using GraphPad Prism (Version 9.4.1) GraphPad Software, San Diego, CA, USA. Negative percent viability values are plotted as zero for the figures shown. Viral titer was calculated by RT-qPCR using cycle quantification (Cq) values that reflect the change in cycles needed to detect viral RNA for the SARS-CoV-2 N1 protein.

## 3. RESULTS

In this paper, the infectivity of the 3 coronavirus variants, wild type (USA/WA1), delta (B.1.617.2), and omicron (B1.1.529), across multiple cell lines were studied. Following infection of VeroE6 cells, supernatants from each replicate at 5 days post-infection were processed to quantify viral titers using RT-qPCR, where lower Cq numbers represent high viral titers and cell viability by LDH release.

As expected, WT SARS-CoV-2 destroyed VeroE6 monolayers at viral doses of 1 PFU to 1000 PFU (Fig.1(c)). WT at a dose as low as 1 PFU resulted in a maximum loss of cell viability while producing a similar amount of viral genome across all infection doses tested (Fig.1(a,b)). Interestingly, for the lowest viral dose of 0.1 PFU, the amount of virus produced and the visual appearance of disrupted monolayers was similar to those at higher PFUs; however, LDH release did not detect this. A careful comparison of LDH results with images reveals a trend of disrupted monolayers at PFU titers lower than the breakpoint titer detected by LDH. For example, at 0.1 PFU, the visually disrupted monolayer did not correspond with a measure of maximum cell viability. In contrast, Omicron and Delta variants failed to reduce host cell viability, and virus titer increased only for Omicron at 1000 PFU. It is of note that Omicron titers were equivalent to WT at 1000 PFU without an appreciable reduction in cell viability. Delta and omicron were unable to cause CPE with any of the infection doses tested except for 1000 PFU for omicron (Fig.1(c)).

**Figure 1.**
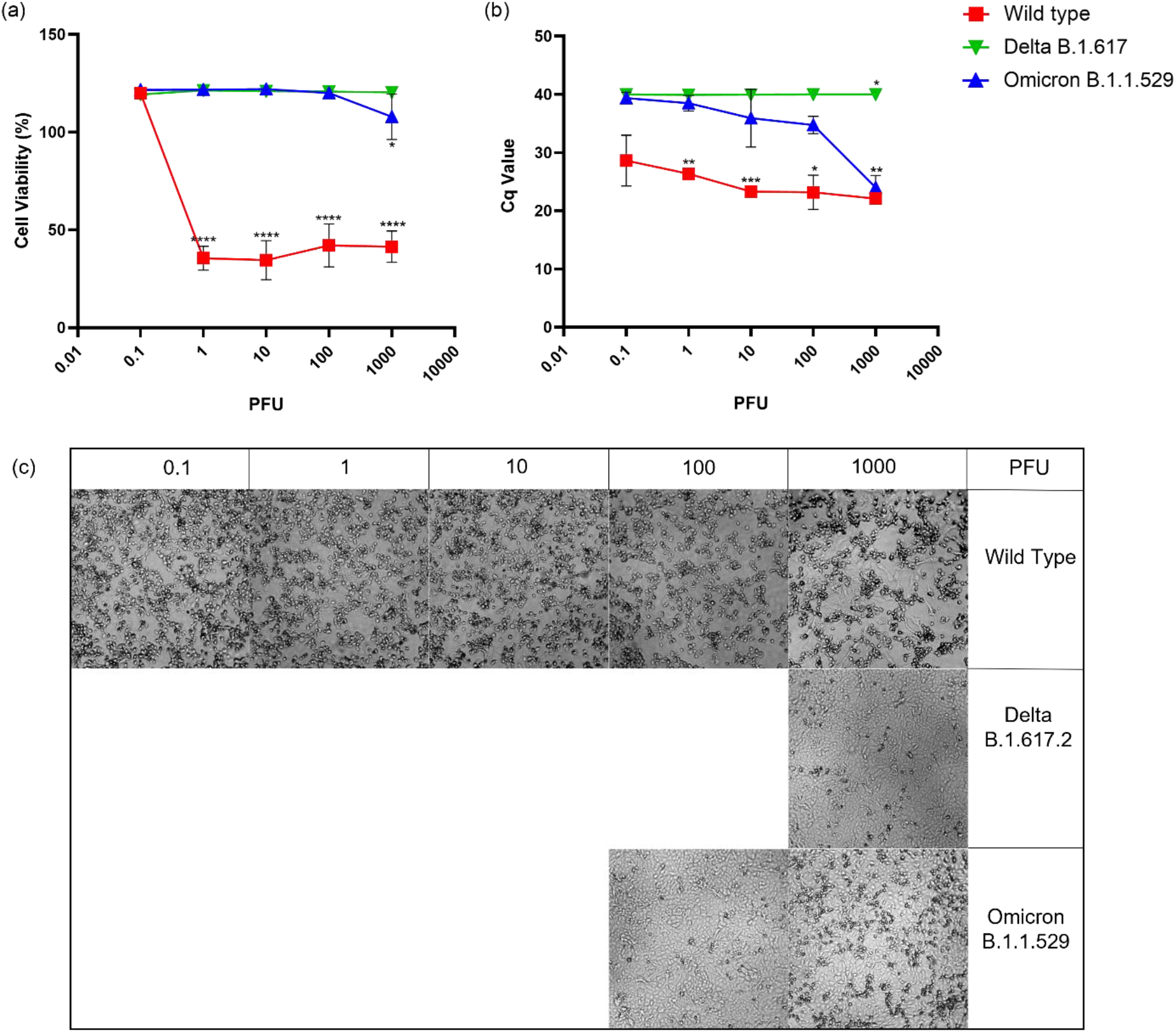
Infection of VeroE6 cells with wild-type, delta, and omicron strains. a) % cell viability of infected cells calculated using LDH released in the supernatant. All values are normalized to uninfected cell controls. b) Cycle quantification (Cq) values measured by RT-qPCR in supernatant from infected cells 5 days post-infection. (c) Brightfield images on low magnification of cells infected with indicated PFU. Images of monolayers bracketing the viability breakpoints are shown. (*p<0.05, **p<0.01, ***<0.001, ****<0.0001)

Employing an identical experimental setup to that of Vero E6 cells, VeroE6 high-expressing endogenous ACE2 (VeroE6/ACE2) were infected with all three variants (Fig.2). For WT SARS-CoV-2, all infection doses reduced cell viability and increased viral titers except 0.1 PFU that did not lower the cell viability in VeroE6/ACE2 (Fig. 2(a,b)). Interestingly, the cells infected with 0.1 PFU of WT showed rounded cells despite the high cell viability (Fig. 2(c)). Delta variant failed to reduce cell viability and increase viral titers at any infection dose except the 1000 PFU, where high viral titers and CPE was observed (Fig. 2). The Omicron variant significantly reduced the cell viability only at the 2 highest viral doses, where CPE was also observed (Fig.2(c)), 100 and 1000 PFU, which corresponded with viral titers produced as well (Fig.2(b)).

**Figure 2.**
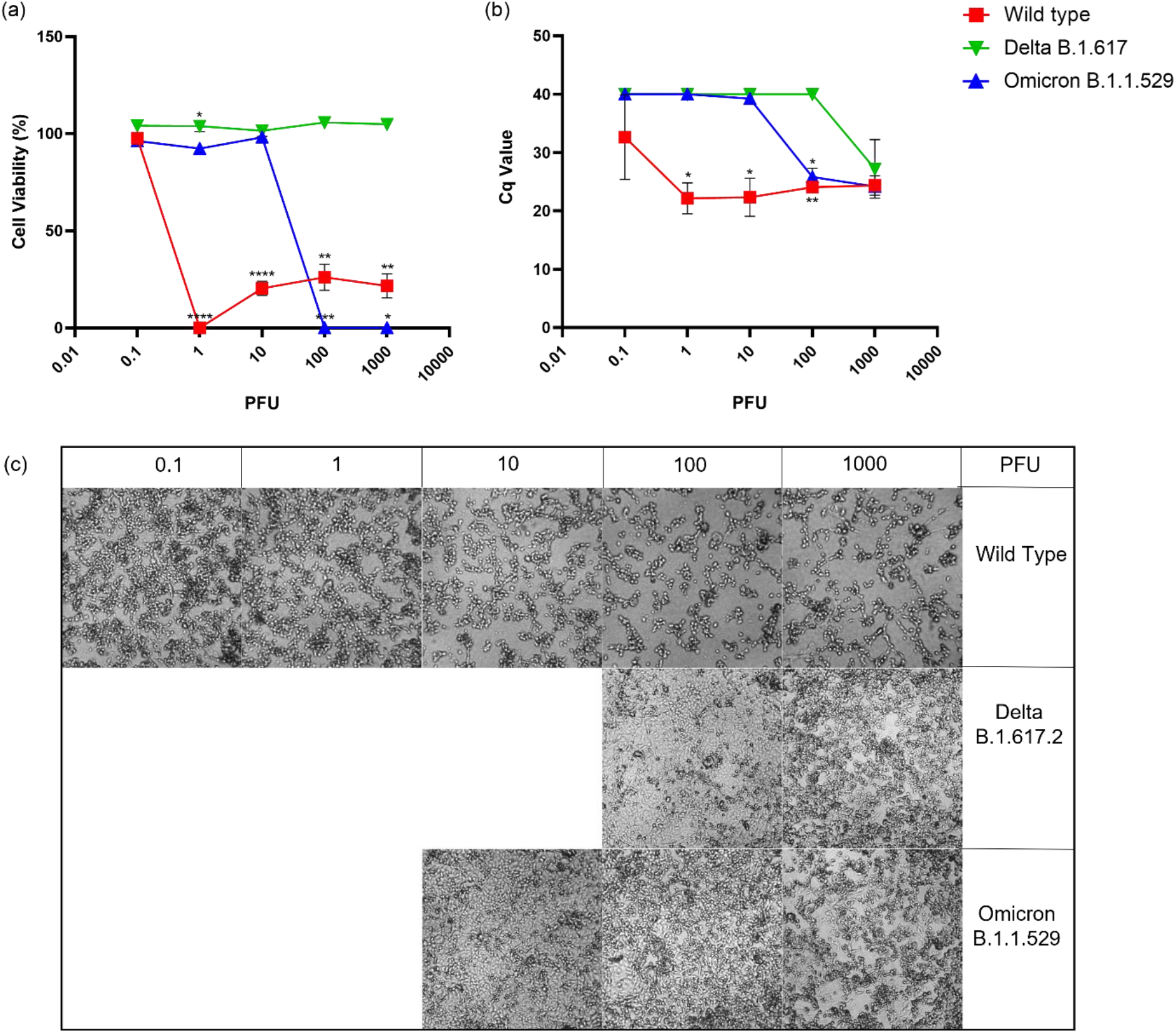
Infection of VeroE6 cells highly expressing endogenous ACE2 with wild-type, delta, and omicron strains. a) % cell viability of infected cells calculated using LDH released in the supernatant. All values are normalized to uninfected cell controls. b) Cycle quantification (Cq) values measured by RT-qPCR in supernatant from infected cells 5 days post-infection. (c) Brightfield images on low magnification of cells infected with indicated PFU. Images of monolayers bracketing the viability breakpoints are shown. (*p<0.05, **p<0.01, ***<0.001, ****<0.0001)

When VeroE6 expressing human ACE2 and TMPRSS2 (VeroE6/hACE2/TMPRSS2) were infected with a viral titration of WT strain, all tested infection doses reduced the cell viability, showed high CPE and high viral titers (Fig.3). On the other hand, both delta and omicron reduced cell viability and had high viral titers at all infection doses except 0.1 PFU (Fig 3(a,b)). CPE caused by omicron infection followed the same trend, where no CPE was observed at the lowest infection dose. In contrast, even with low cell viability and high viral titers at 1 PFU of the delta variant, CPE was not observed (Fig. 3(c)).

**Figure 3.**
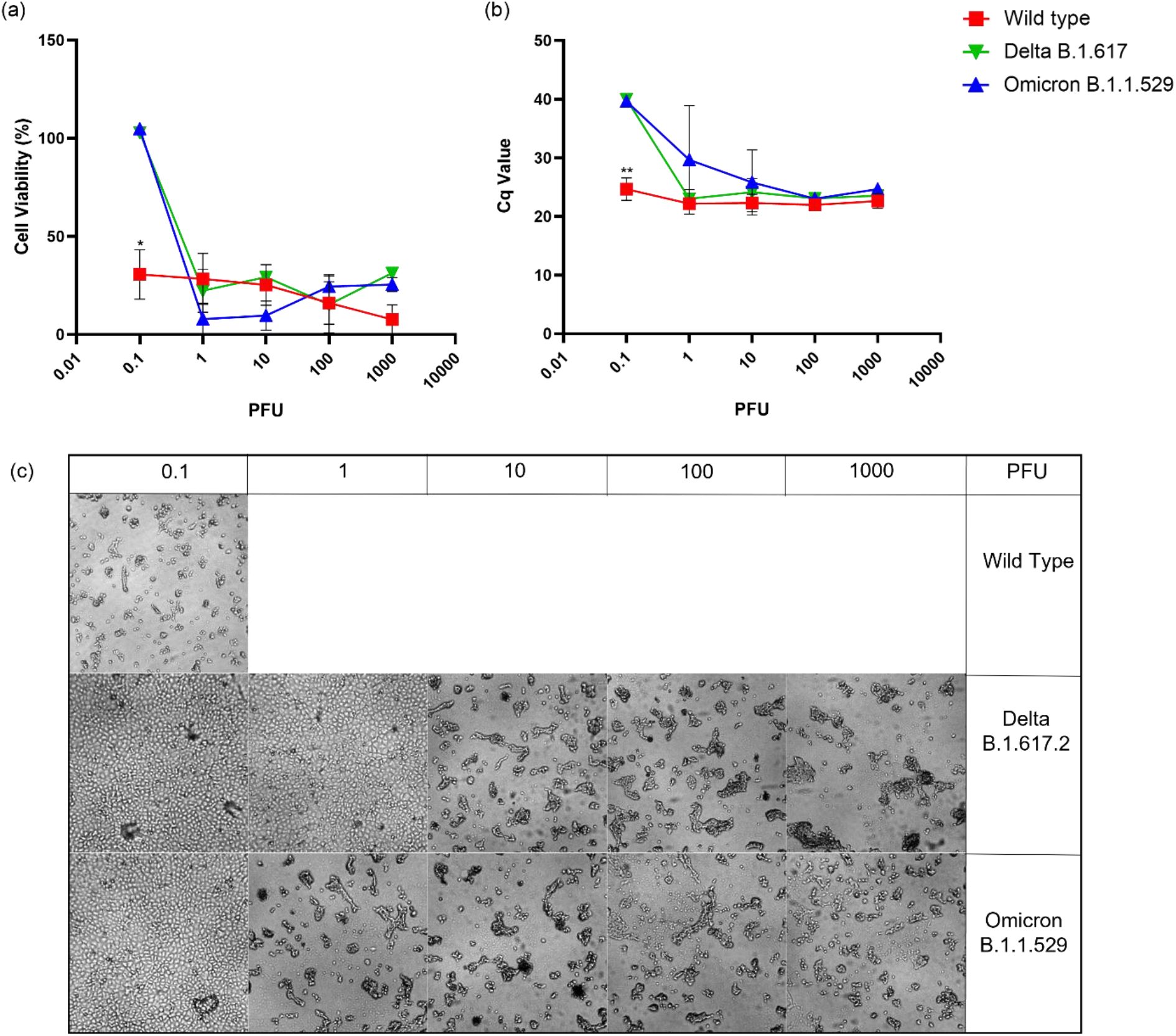
Infection of VeroE6 cells highly expressing human ACE2 and TMPRSS2 with wild-type, delta, and omicron strains. a) % cell viability of infected cells calculated using LDH released in the supernatant. All values are normalized to uninfected cell controls. b) Cycle quantification (Cq) values measured by RT-qPCR in supernatant from infected cells 5 days post-infection. (c) Brightfield images on low magnification of cells infected with indicated PFU. Images of monolayers bracketing the viability breakpoints are shown. (*p<0.05, **p<0.01)

Human lung cell lines, Calu3 high expressing ACE2 (Clu3/hACE2) and A549 (Fig. 4), were infected with the three SARS-CoV-2 variants. No significant reduction in cell viability was measured (Fig. 4(a,c)) at any infection dose for all three variants. Viral titers measured after RT-qPCR were dose dependent for WT and omicron infected lung cells. Specifically for omicron, an increase in viral titers was observed at infection dose 10 PFU. For cells infected with delta, low viral titers were observed at all infection doses (Fig. 4(b,d)) in both cell lines.

**Figure 4.**
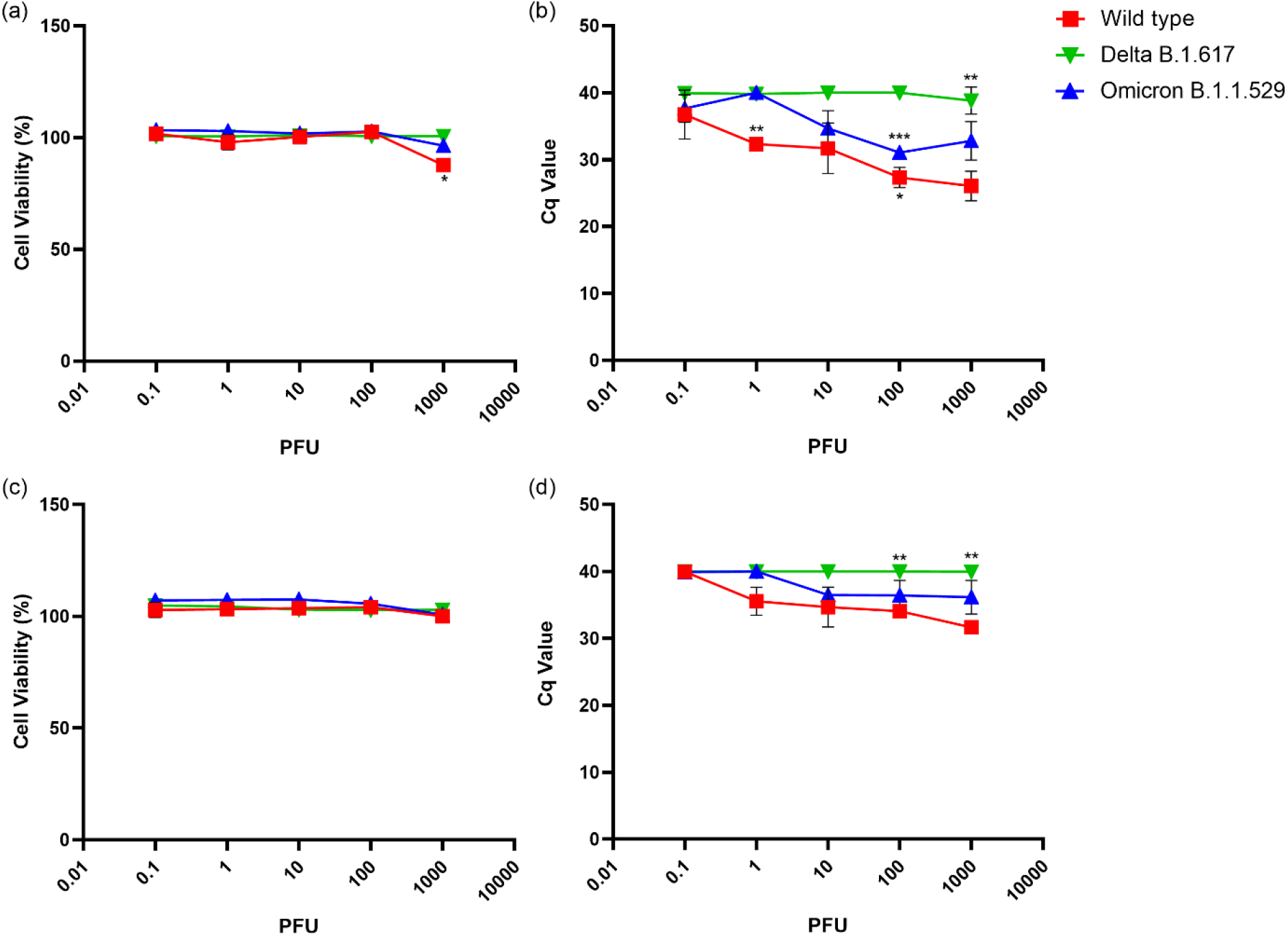
Infection of Calu3 cells highly expressing human ACE2 (Calu3/hACE2) and A549 cells with wild-type, delta, and omicron strains. a) and c) % cell viability of infected Calu3/hACE2 and A549 cells calculated using LDH released in the supernatant, respectively. All values are normalized to uninfected cell controls. b) and d) Cycle quantification (Cq) values measured by RT-qPCR in supernatant from infected Calu3/hACE2 and A549 cells 5 days post-infection, respectively. (*p<0.05, **p<0.01, ***<0.001)

## 4. DISCUSSION

There is extensive ongoing research on the different variants of SARS-CoV-2 that have emerged in the last three years. Each variant with unique infection kinetics and pathogenicity is investigated in numerous different models. Having a defined cell culture model for all the variants is critical for invitro studies, such as neutralization assays or antiviral testing. Understanding the infection kinetics of different variants in the same environment will not only give us a better understanding of the viral pathogenesis but also help us choose the right infection model for in vitro experiments.

We have highlighted two important parameters to consider when selecting a cell culture model; cytopathic effect (CPE) and replication. During viral infection, the contents of the virus are transferred into the cytoplasm of the cell, where the host cell machinery can be used for viral replication. After cell entry, the viral particles produced are released through lysosomal exocytosis into the extracellular matrix (19). SARS-CoV-2 infected cells have Spike protein on their surface that can bind to the ACE2 receptor on adjacent uninfected cells that can cause cell-cell fusion called syncytia (20). Syncytia formation, along with apoptosis, autophagy, necroptosis, and inflammation activation in the host cells, can induce cell death or cytopathic effect (CPE) in SARS-CoV-2 infected cells (21,22). CPE is measured by measuring the cell viability of the infected cells using lactate dehydrogenase (LDH) released in the media after a cell has succumbed to an infection. CPE can be directly correlated to the amount of LDH released in the cell culture media, which can be corroborated visually under a microscope. Cells that undergo CPE start rounding up and die, leaving holes in the monolayer. SARS-CoV-2 virions released in the supernatant of infected cells were quantified using RT-qPCR. The increase in RNA copies (inversely related to Cq value) in the supernatant correlates with viral replication.

Both omicron and delta replication was attenuated compared to wild type in VeroE6 cells at 120 hours post-infection (Fig. 1(b)), which was also observed by Shuai et al. at 24- and 48 hours post-infection (23). At 72hr post-infection, Zhao et al. observed omicron and delta had comparable viral replication in VeroE6 cells at 0.1 PFU infection dose (24), which was also observed at 120hour post-infection in this study(Fig. 1(b)). However, when the infection dose is increased to 10 - 1000 PFU, omicron shows higher replication than delta in VeroE6 cells (Fig. 1(b)). Both omicron and delta failed to cause a reduction in cell viability of VeroE6 cells at all the infection doses tested, except 1000 PFU for omicron, which produced a noticeable CPE when observed under the microscope(Fig.1(c)). This attenuated infectivity of omicron can be attributed to its decreased fusogenicity and pathogenicity (23,25). At the same time, the reduced infectivity of delta can be due to its dependence on TMPRSS2 for cell entry (24,26) which is expressed marginally in VeroE6 cells (17).

When ACE2 is overexpressed in VeroE6 cells, the omicron variant Cq value is reduced from 38 in VeroE6 to 25 in VeroE6/ACE2. This increase in replication was not observed for delta (Fig. 1 and Fig. 2). Similarly, with the higher expression of ACE2 in Calu3 cells, viral release in the supernatant of Calu3/hACE2 cells infected with omicron strain was higher than that of delta at day 5 post-infection (Fig. 4), compared to Calu3 cells where omicron has been shown to release fewer viral particles than delta at 2 days post-infection (23–25). In both Calu3 and VeroE6, overexpression of ACE2 increased the infectivity of omicron, suggesting an increased importance of ACE2 for the pathogenesis of omicron compared to delta. This could be due to omicron spike protein having a higher binding affinity to ACE2 than delta spike protein, as shown in previous studies (27–30). Even though omicron spike protein binding affinity is higher than WT, the inherent susceptibility of VeroE6 cells to WT (Fig. 1) is responsible for the high infectivity of WT in VeroE6/ACE2 cells. At 0.1 PFU for WT strain in VeroE6 and VeroE6/ACE2 cells (Fig. 1 and 2), the % cell viability measured does not correlate with the CPE observed under the microscope. The appearance of monolayer disruption without a release of the cytosolic LDH would suggest that cells lift off the surface prior to the breakdown of the plasma membrane.

Replication of omicron in VeroE6/ACE2 cells was higher than delta(Fig. 2), but when TMPRSS2 is overexpressed, replication of both the variants is comparable (Fig.3). This suggests that the presence of TMPRSS2 has a larger impact on delta replication than it does on omicron replication. Consistent with previous studies (23–25), omicron utilizes TMPRSS2 less efficiently for S cleavage than delta or WT and uses the endocytic pathway instead for cell entry (31).

At 120 hours post-infection, the WT strain shows dose-dependent CPE in VeroE6 but no visible CPE in Calu3/hACE2 or A549s (Fig. 4) (32). Omicron and WT viral titers in the supernatants of Calu3/hACE2 cells were significantly higher than delta. Cell lysates of Calu3/hACE2 have shown higher viral titers for delta compared to omicron (33), suggesting altered viral release of delta from Calu3/hACE2. Similarly, in A549, no CPE was observed at day 5 post-infection for all three variants, with minimal viral release observed for WT and omicron (Fig. 4). Concurrent with other findings, ACE2 and TMPRSS2 expression in A549 cells is less than in other cell lines (34,35) contributing to the low infectivity of A549 cells.

In conclusion, for CPE or plaque-based assays, such as neutralization assays or antiviral testing assays, choosing VeroE6/hACE22/TMPRSS2 would be beneficial over VeroE6 given the formers’ higher sensitivity to SARS-CoV-2 infection. This is a unique study with a parallel comparison of multiple variants across several cell lines. This study highlights the importance of understanding cell line sensitivities to different variants, contributing to future experimental design and understanding the SARS-CoV-2 variant pathogenesis.

## 6. AUTHOR CONTRIBUTIONS

KSP performed scientific experiments, results analysis, first draft, tables, and figures conceptualization. KSP and NBAH maintained cell lines. KSP and BHB wrote the first draft. All authors contributed to manuscript editing and revision and approved the submitted version. All authors contributed to the article and approved the submitted version.

